# A Random Matrix Approach to Single Cell RNA-seq Analysis

**DOI:** 10.1101/2023.06.28.546922

**Authors:** Sivan Leviyang

## Abstract

Single cell RNA-seq (scRNAseq) workflows typically start with a raw expression matrix and end with the clustering of sampled cells. Viewed broadly, scRNAseq is a signal processing workflow that takes a transcriptional signal as input and outputs a cell clustering. Currently, we lack a quantitative framework through which to describe the input signal and assess the dependence of correct clustering on the signal properties. As a result, fundamental questions regarding the resolution of scRNAseq remain unanswered and experimentalists have little guidance in determining whether a hypothesized cell type will be clustered by a particular scRNAseq experiment.

In this work, we define the notion of a transcriptional signal associated with a gene module, show that the tools of random matrix theory can be used to characterize the signal as it moves through a common (PCA-based) scRNAseq workflow, and develop estimates for cell clustering based on the signal properties and, in particular, the signal strength. We give a formula - that can be computed from expression data - for the signal strength, providing a framework through which scRNAseq resolution can be investigated.

## 1 Introduction

Single cell RNAseq (scRNAseq) is often used for de-novo identification of cell types and states. For example, the Human Cell Atlas aims (among other goals) to compile scRNAseq datasets for the exhaustive identification of human cell types [35]. Similarly motivated atlases have been constructed across a range of organisms, tissues, and biological states [34]. Central to the construction of cell atlases, and the application of scRNAseq in general, are the algorithms used in scRNAseq workflows to cluster the sampled cells. Seurat and scanpy are two popular software packages that provide workflows starting with a raw scRNAseq transcription matrix and ending with cell clusters [37, 1]. Within these packages, the workflow can take different forms, but roughly a subset of genes is selected, the resultant expression matrix is normalized and scaled, the scaled expressions are embedded in a low dimensional Euclidean space, a cell graph based on the embedding is formed, and cell clustering is performed on the graph.

Since RNAseq captures only the transcriptional state of a cell, cell clustering reflects differential expression of genes across different cell types or states. If we view the differential expression patterns as a signal, we can ask how different expression levels in cell states must be for the cell states to be properly clustered? Or, put another way, what is the resolution of the scRNAseq workflow? Over the past decade, tools and analyses have been developed for scRNAseq normalization (e.g. [16, 21, 10, 15]), dimension reduction (e.g. [40, 6, 24]), and graph construction (e.g. [42, 22, 17]), but we lack a quantitative framework through which to define a transcriptional signal and analyze it as it moves through the scRNAseq workflow,. As a result, experimentalists have little guidance in assessing whether a particular scRNAseq experiment will reveal a hypothesized cell type and fundamental questions regarding the resolution of scRNAseq remain unanswered.

A paradigm of transcription analysis has been the existence of gene modules, broadly defined as groups of genes with expression levels that vary in a correlated manner across cell states [14, 33, 5, 44, 36]. In this work, we assume a statistical model for a gene module, define the module’s signal and signal strength, and then exploit existing results in random matrix theory (RMT) to analyze clustering as signal strength varies. As an example, suppose scRNAseq is performed on a collection of cell types, say PBMCs. Within those cell types, suppose there is a particular cell type, say B cells, and a collection of genes that are differentially expressed in the B cells relative to the other cell types. Those differentially expressed genes form our gene module and we study the clustering of B cells as a function of the gene module’s signal strength.

RMT has been used in scRNAseq analysis by Aparicio et al. [2]. The authors showed that scRNAseq expression matrices are approximated by classical RMT results, but that some deviations occur due to sparsity. They developed a gene selection and normalization algorithm by exploiting these deviations. Here, we work downstream in the scRNAseq workflow. We define the signal in the expression matrix and follow the signal through dimension reduction by PCA and graph construction by k-nearest neighbors (knn).

RMT results provide explicit formulas for the PCA under the so-celled spiked model [18, 19], which decomposes a matrix into a sum of a deterministic matrix - the spike - and a random matrix. Our statistical model of a gene module allows us to decompose the scaled expression matrix into a sum of a spike, which encodes the signal, and a random matrix, which encodes noise. We apply RMT to develop formulas describing the signal as it passes through the PCA and graph construction. As an endpoint, our formulas allow us to predict the fraction of cells that have the same cell state as their nearest neighbor in the knn graph, which allows us to quantify clustering.

Our formulas are approximations because gene modules do not exactly obey our statistical model, because we assume a particularly simple form for the noise of scRNAseq and because the RMT results we use assume a large sample limit. We examine the accuracy of our approximations by comparing their predictions to the true values generated by running the scRNAseq workflow on simulated and real datasets. At least for the datasets we consider, our approximations are relatively accurate.

Our results provide a quantitative framework through which scRNAseq resolution can be understood. We characterize signal strength in terms of the signal-to-noise ratios (SNR) of genes and we provide a biologically interpretable formula for a gene’s SNR. The SNR of a gene can be estimated from scRNAseq data, allowing for estimates of the signal strength and prediction of downstream clustering.

## 2 Results

### Statistical Model of the Expression Matrix

The scRNaseq workflow starts with a (raw) expression count matrix *X* for which we define a statistical model. Let *X* be a *C* × *G* matrix with *X*_*cg*_ the expression count of gene *g* in cell *c*. We split the *G* genes into *G*^base^ baseline genes and *G*^pert^ perturbation genes and write,

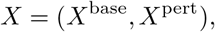

where *X*^base^ and *X*^pert^ are restricted to the baseline and perturbation genes, respectively.

We assume that the perturbation genes are differentially expressed across two cell states, which we call the perturbation states and label as *S*_1_ and *S*_2_. Letting the fraction of sampled cells in the two respective states be *p*_1_ and *p*_2_, we assume that each cell is in one of these two states, making *p*_1_ + *p*_2_ = 1. The sampled expression counts are distributed as

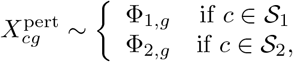

where Φ_1,*g*_, Φ_2,*g*_ are random variables. In most statistical models of gene expression, the Φ are chosen as negative binomial distributions. For example, Φ_1,*g*_ = NB(*µ*_1,*g*_, *θ*) and Φ_2,*g*_ = NB(*µ*_2,*g*_, *θ*), where *µ*_1,*g*_ and *µ*_2,*g*_ would be the mean expression counts for gene *g* in the two cell states, respectively,, and *θ* is a shared dispersion. However, the distribution assumed is not an essential part of our analysis. What is essential to our analysis is that 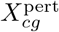 is drawn independently for each index pair *c, g*. This means that the correlation in expression counts between two cells is entirely determined by the cell states, i.e. *S*_1_ and *S*_2_.

*X*^base^ is assumed to be composed of an arbitrary number of gene modules, which we do not explicitly model. Baseline genes may be homogeneously expressed across the cells, which corresponds to a gene module with Φ_1,*g*_ ∼ Φ_2,*g*_.

To provide concreteness, imagine the expression matrix *X* samples PBMCs and that the perturbation genes are a particular collection of genes differentially expressed in B cells relative to the other cell types. Then the two cell states, *S*_1_ and *S*_2_, indicate whether a cell is a B cell or not. Importantly, the baseline expression matrix *X*^base^ can include genes with any differential expression pattern: there is no restriction that the differential expression pattern not somehow involve B cells. We use the model to compare the clustering of B cells - or more generically the clustering of cells in states *S* 1 and *S* 2 - with and without the perturbation genes included in the expression matrix.

Given a count matrix *X*, scRNAseq workflows typically include quality control (QC) and gene selection steps that filter the sampled cells and genes. We assume that *X* represents the post-QC and gene selection matrix. Subsequent to these pre-processing steps, scRNAseq workflows construct a normalized count matrix *Y* and then a scaled count matrix *Z*. Normalization approaches vary; we normalize by adjusting for library size and then log-transforming, as implemented in Seurat’s NormalizeData function. Defining 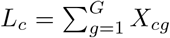, the library size of cell *c*, and *L* = 10000,

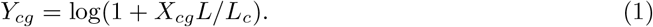

The *Z* matrix is formed by centering each column of *Y* (i.e. each gene) and then scaling the column to Euclidean length 1 As with *X, Z* is decomposed into *Z* = (*Z*^base^, *Z*^pert^).

### RMT Decomposition of the Scaled Expression Matrix

The assumptions of our statistical model provide the following decomposition of the perturbation portion of the scaled expression matrix *Z*,

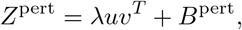

where λ is a scalar, *u* and *v* are vectors of dimension *C* and *G*^pert^, respectively. *uv*^*T*^ is the outer product forming a *C* × *G*^pert^ matrix and *B*^pert^ is a *C* × *G*^pert^ random matrix. Importantly, the mean value of each entry of *B*^pert^ is zero, and so *B*^pert^ can be viewed as noise. Following RMT terminology, we refer to the matrices *λuv*^*T*^ and *B*^pert^ as the perturbation spike and bulk matrix, respectively.

Formulas for λ, *u* and *v* provide a biological interpretation of the spike. We define λ^2^ as the signal strength of the perturbation module. Letting *g* range over the columns (i.e. perturbation genes) of *Z*^pert^,

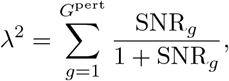

where SNR_*g*_ is the signal to noise ratio of gene *g* and is given by,

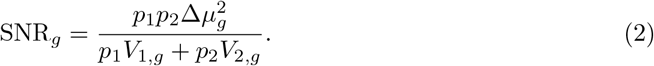

Above, Δ*µ*_*g*_ is the difference in the mean normalized gene expressions in the two cell states for gene *g*,

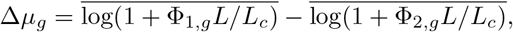

where the bars above represent mean value over the distribution of Φ. Δ*µ*_*g*_ is the commonly used logFC metric for differential expression. *V*_1,*g*_ and *V*_2,*g*_ are the variances in gene expression within the two cell states for gene *g*, i.e. *V*_1,*g*_ is the variance of *Y*_*cg*_ when *c* is assumed to have cell state *S*_1_. In the typical negative binomial model of gene expression, these variances are simply the variance of a negative binomial with fixed parameters. The SNR is the ratio of variance across the cell module states to the variance within cell module state. Importantly, if the cell states are known we can estimate SNR_*g*_ from data using standard mean and variance estimators.

We define the vector *u* in the spike matrix as the module signal. *u* is a scaled version of the cell state indicator function and therefore encodes the module states across the cells. (To construct *u* set coordinate *c* of *u* equal to 1 if cell *c* is in state *S*_1_ and 0 otherwise, center *u* to mean 0, and scale *u* to length 1.) Finally *v* simply collects the λ_*g*_: *v*_*g*_ = λ_*g*_/λ.

Following on our assumption that *X*^base^ is formed by multiple modules,

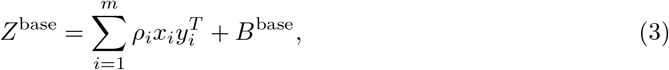

where *m* is the number of modules, the summation expression is a deterministic matrix and *B*^base^ is a mean zero, random matrix. Importantly, (3) is what we need to assume about *Z*^base^. The assumption that *X*^base^ is formed from multiple gene modules gives biological intuition, but is more restrictive than necessary. We call the respective summands in (3) the baseline spike and baseline bulk matrix. The 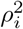 represent the signal strength of the baseline modules. Finally, combining the expressions for *Z*^pert^ and *Z*^base^,

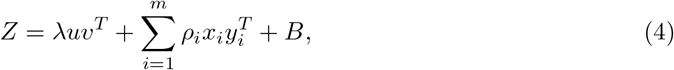

where we call *B* = (*B*^base^, *B*^pert^) the bulk matrix. (Above, we extend *v* and *y*_*i*_ to all *G* genes by setting to 0 all coordinates corresponding to baseline and perturbation genes, respectively.)

### The RMT Approximation

The main mathematical claim of this paper is that we can use RMT results to analyze the scRNAseq workflow on *Z* when we replace *B* by a matrix *N* that meets standard RMT assumptions. The simplest choice for *N* would be a matrix of independent normally distributed entries with the columns scaled to sum to 1. However, this ignores the role of the spikes. Since the columns of *Z* are scaled to norm 1, the presence of spikes with large λ or *ρ*_*i*_ typically leads to *B* with column norms less than 1. We could use the column norms of *B* to define *N*, but we want *N* to be independent of the perturbation module. With this in mind, we define *N* with column norms from the baseline bulk. Let 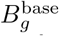 be the *g*th column of the baseline bulk, and define *χ* as the random variable with value 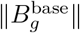 where *g* is uniformly chosen from the *G*^base^ genes. Then we define *N* by

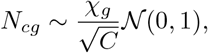

where *χ*_*g*_ is independently sampled from *χ* for each column *g. N* satisfies the assumptions in most RMT papers, e.g. [25, 18, 3, 4], and in particular the results of Benaych-Georges and Nadakuditi [8, 7], which underlie most of our formulas below. Using *N*, we define 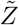 as *Z* in (4), but with *N* replacing *B*,

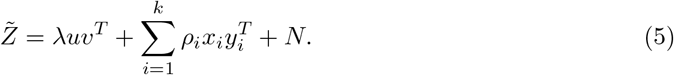

We use 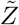 to develop formulas describing the dimension reduction and graph construction steps of the scRNAseq workflow and then examine the accuracy of these formulas by running the scRNAseq workflow on simulated and real datasets.

### Analysis of PCA and Graph Construction

The analysis of 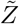 depends on the relationship between the module spike (*λuv*^*T*^) and the baseline spike 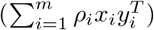 in (4). Our analysis increases in complexity as we move sequentially through the following cases.

*1. Homogeneous Baseline* : The baseline spike is absent, meaning 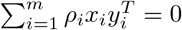

*2. Uncorrelated Baseline* : The baseline spike can be present but *u* and the *x*_*i*_ are not correlated.

*3. General Baseline*: The baseline spike can have any form.

These cases have biological significance. To see how, take *u* to be simply 1 or 0 based on whether a cell is in state *S*_1_ and *S*_2_ and imagine that the *x*_*i*_ represent similar indicator functions for given cell types. For example, if we have a collection of PBMCs, then *x*_1_ indicates if a cell is a macrophage, *x*_2_ indicates if a cell is a T cell, etc. A homogeneous baseline means we have no *x*_*i*_, which might be the case for a homogeneous cell line, e.g. all the cells are macrophages. A uncorrelated baseline means that knowing the cell type tells us nothing about the perturbation state. For example, if the two perturbation states are a control and a stimulated state, an uncorrelated baseline would be the case if all cell types respond similarly (in terms of transcription) to the stimulation.

We first present our analysis for the case of a homogenous baseline which gives particularly simple and instructive formulas. Let 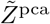 be the *C* × *k* PCA dimension reduction of the scaled expression matrix 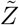, where *k* is the number of principal components used. In this case, 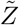 forms a spiked RMT model with a single spike and the celebrated BBP phase transition of RMT [3, 32, 8] establishes (in the large sample limit) a threshold *τ* below which the signal is lost and above which the signal is present in the dominant principal component. Theorems 2.9 and 2.10 in [8] give the following form for the columns of 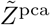,

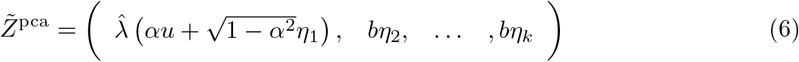

where *b* is the maximum eigenvalue (singular value) of *N* and the *η*_*i*_ are independent *C* dimensional random vectors with independent coordinates distributed as 𝒩 (0, 1*/C*). The dimension reduction step acts as a filter, taking<u>as inpu</u>t the module strength and signal, *λ* and *u*, and giving an output of 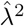 and 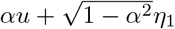. The variables 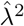 and *α* are functions of the signal strength *λ*^2^ and depend on *N*, with formulas derived in [8] and described in the Materials and Methods.

Figure 1 shows the value of 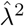 and *α*^2^ as λ^2^ varies for the Kang et al. dataset (described below). When *λ < τ*, *α* = 0 and 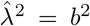. In this case no portion of *u* is captured by the dimension reduction and the signal has been completely lost. As λ increases beyond *τ*, *α* increases and *u* is increasingly captured in the dimension reduction. Importantly, the signal is always restricted to the dominant principal component.

**Figure 1.**
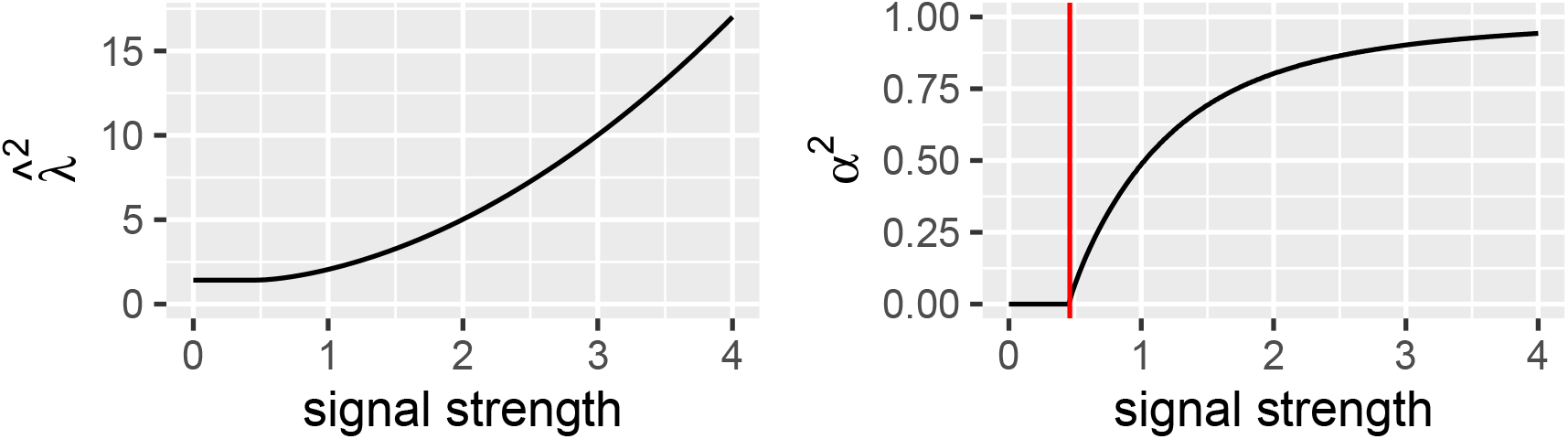
The PCA Acts as a Filter on the Module Signal. The module strength and signal λ^2^ and *u*, respectively, are filtered by the PCA dimension reduction to 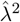 (left panel) and 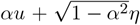(*α*^2^ shown in right panel), respectively. The coordinates of *η* are normally distributed and represent noise. Values shown are for Kang et al. dataset. The module signal is completely lost for *λ < τ* (here *τ* = 0.46) but is recovered as λ increases beyond *τ*.

After dimension reduction, the variance of the data in the direction of the signal *u* is given by 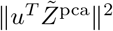 and we define this quantity as the filtered signal strength. Ignoring terms of order 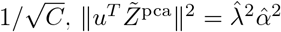.

The next step in the workflow involves construction of a cell graph based on the distances between cells (rows) in the dimension reduction matrix 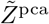. Graph algorithms vary; k nearest neighbor (knn) and UMAP graphs ([26]) are common choices. Our analysis assumes a knn graph (but see below for connections to UMAP). To analyze the graph-based clustering, let nn(*G*) be the fraction of cells in state *S*_1_ whose nearest neighbor is also in state *S*_1_, where *G* is the knn graph.

Let 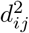 be the distance under the dimension reduction between the cell *i* and *j*. Using (6) allows us to write down a formula for 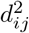 and some simplification (see Materials and Methods) gives,

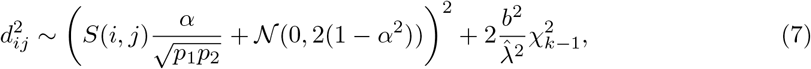

where *S*(*i, j*) is 0 if cells *i* and *j* share the same perturbation state and 1 otherwise, and 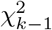 is a chi-squared random variable with *k* − 1 degress of freedom. (7) is instructive. The term 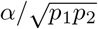 captures the module signal, while the chi-squared term represents noise. As the module’s signal strength λ^2^ increases, *α* will increase to 1 eventually separating cells in different states by a squared distance 1*/p*_1_*p*_2_. Further, as the λ increase, so will 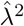 and this will dampen the noise term.

Assuming that *d*_*ij*_ is independent over *j* for a fixed cell *i*, we generate samples for *d*_*ij*_ over all cells *j* ≠ *i*, and compute the expected value of nn(*G*) using a Monte Carlo approximation.

The assumption of a homogeneous baseline leads to a simple PCA, with all principle components being normally distributed except for the first. If we assume an uncorrelated baseline, the PCA-filtered signal is still contained in a single principle component - not necessarily the first - but the other principle components reflect the structure of the baseline expression and need not be normally distributed. If we allow the baseline spike to be correlated with the signal, then the PCA-filtered signal can be spread across multiple principle components. Importantly, in this setting, the BBP transition can disappear, meaning that some portion of the signal will be captured in the PCA filtering regardless of the signal strength. See the Materials and Methods for mathematical details for the uncorrelated and general baseline cases.

### Model Evaluation

To evaluate the accuracy of our predictions, we focus on two metrics: the strength of the signal after PCA dimension reduction (filtered signal strength) and the fraction of cells that have the same perturbation state as their nearest neighbor in the knn graph (nn(*G*)). We ran the scRNAseq workflow on a simulated dataset generated by splatter [43] and the real datasets of Kang et al [20], Mead et al. [27], Ordovas-Montanes et al. [31], and Zheng et al [45] and compared the true and predicted values for the two metrics.

For all datasets, we started with a raw count matrix *X* and applied QC and gene selection for the 1000 most variable genes. We then identified a subset of these 1000 genes to serve as the perturbation genes; all other genes were baseline genes. The choice of the perturbation genes was dataset specific. We constructed *X*^*𝓁*^ = (*X*^base^, *X*^pert,*𝓁*^) where *X*^pert,*𝓁*^ is the perturbation expression matrix *X*^pert^ restricted to the first *𝓁* perturbation genes. Importantly, the baseline stayed fixed as *𝓁*, the number of perturbation genes included in *X*, varied. As *𝓁* increases, the signal strength λ^2^ of the perturbation module increases and we investigated the accuracy of our predictions as a function of λ^2^. The perturbation genes were differentially expressed across two cell states, *S*_1_ and *S*_2_, which were dataset specific. Importantly, in our datasets we knew the state of each cell, so we could compute the SNR of each perturbation gene and consequently the signal strength.

To ground this general description, we describe its implementation for the Kang et al dataset. The Kang et al. dataset is composed of untreated and interferon *β* (IFN) stimulated PBMC from 8 lupus patients. Kang et al annotated the sampled cells into 8 cell types (B cells, NK cells, etc.) in IFN stimulated and control states. We considered the full Kang dataset as well as the dataset restricted to individual cell types. For simplicity, here we describe the datasets formed from individual cell types. For the given cell type, we identified genes that were differentially expressed between IFN stimulated and control cells. The differentially expressed genes, referred to as interferon stimulated genes (ISGs) [38], were the perturbation genes and the two perturbation states, *S*_1_ and *S*_2_, correspond to IFN stimulated and control states. We then investigated clustering of IFN stimulated cells as the signal strength of the perturbation module, which in this case we refer to as the ISG module, varies.

### Simulated Dataset

We used the splatter R package [43] to simulate a dataset analogous to the Kang et al dataset. The simulated dataset contained 5 cell types, split into IFN and control states. Perturbation genes were analogous to ISGs. The signal strength was varied by changing the number of ISGs included in the expression matrix. Figure 2A shows UMAPs generated for different signal strengths, corresponding to different number of ISGs. When the signal is 0, clustering corresponds to the 5 cell types and no clustering based on IFN stimulation occurs. As the signal is raised, clustering associated with IFN stimulation occurs. Figures 2B,C compare our predictions for the filtered signal strength and the nearest neighbor metric to true values. The fits are excellent.

**Figure 2.**
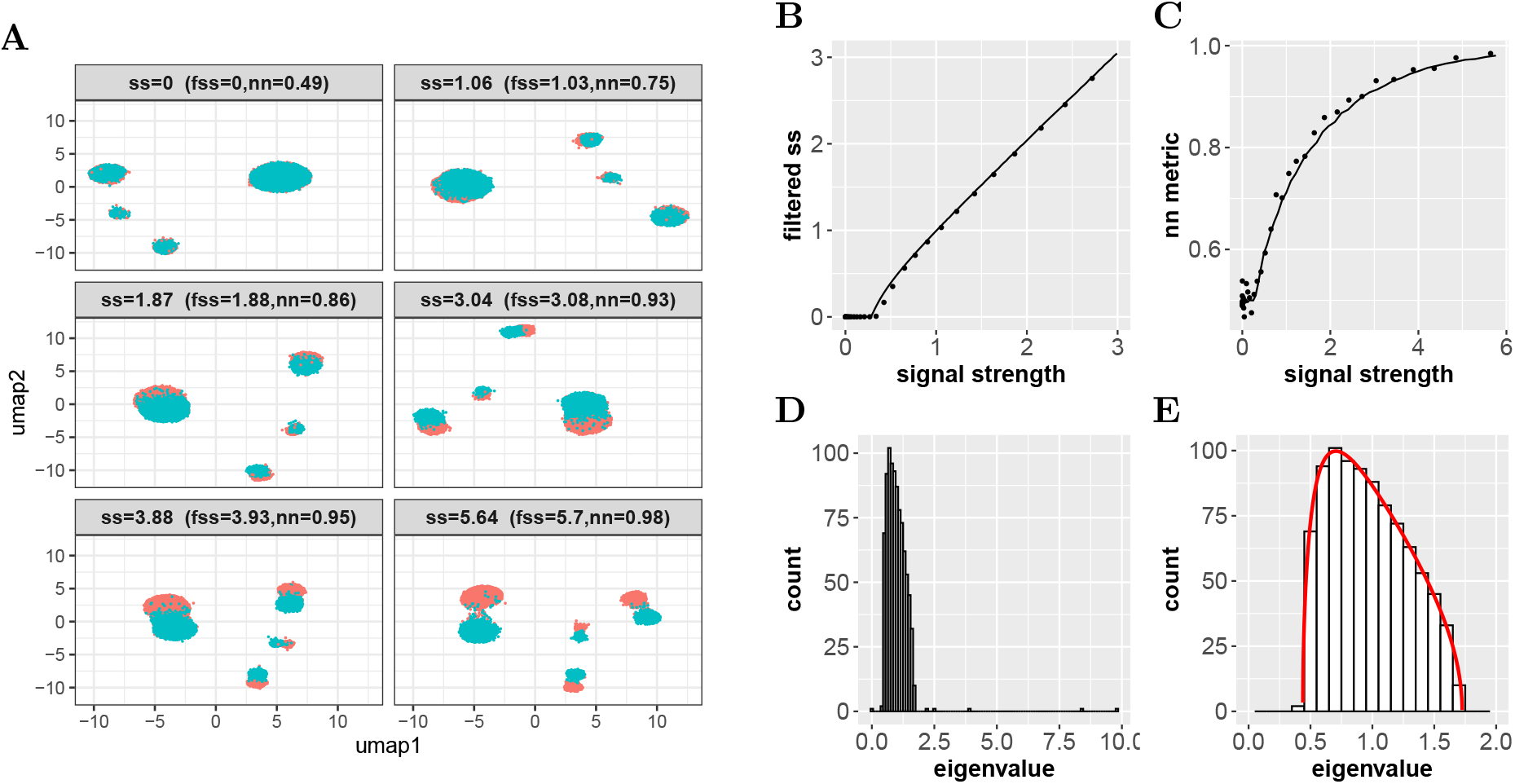
Simulated Dataset. (A) UMAPs generated by Seurat for the simulated dataset. Above each UMAP are the signal strength (ss), the filtered signal strength after PCA (fss), and the nearest neighbor metric on the knn graph (nn). At signal strength 0, the five cell types are clustered. As signal strength increases, IFN stimulated (red) and control cell (blue) states become clustered. (B,C) True (dots) and predicted (lines) values for the filtered signal strength (panel B) and the nearest neighbor metric (panel C) versus the signal strength. (D) Shown are the eigenvalues of the scaled expression matrix. Isolated eigenvalues are the perturbation and baseline spike strengths (λ^2^ and 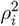 in (4)) and the grouped eigenvalues are from the bulk matrix. (E) A zoom in of the bulk in panel D. The red line represents the eigenvalues of our approximation matrix *N* that satisfies the assumptions of RMT.

Plainly visible in the filtered signal strength is a BBP threshold (roughly 0.5 for this dataset) below which no signal passes through the PCA. The nearest neighbor metric corresponds well to the clustering seen in the UMAPs. When the signal strenght is 0, clustering of cells based on IFN stimulation is completely random, leading to a nearest neighbor metric of 0.5 since cells were equally split between IFN stimulation and control. When the signal strength is above 5, the UMAP shows almost complete clustering by IFN stimulation state and the nearest neighbor metric is almost 1.

Our predictions are based on replacement of the true bulk matrix in the scaled expression matrix by a bulk matrix, *N*, satisfying RMT assumptions. For the simulated dataset, we can decompose the scaled expression matrix into its spike and bulk components and compare the true bulk to the approximate bulk. Figures 2D,E show the decompostion of the scaled experession matrix and the fit of *N* to the true bulk through the matrix eigenvalues. The eigenvalues of *N* are an excellect fit for the eigenvalues of the bulk.

### Single Celltypes

We next consider individual cell types within the Kang et al datasets. The baseline expression of a single cell type has less structure than expression across multiple cell types, making these datasets a middle ground between simulated datasets and more complex, real datasets. Further, subsampling from the same dataset allowed us to compare results across subsamples without convolving batch effects. Each cell type in the Kang et al dataset is roughly split between IFN stimulated and control cells. To investigate the accuracy of our predictions when cell frequencies are skewed, we also subsampled cells within each cell type to make the IFN stimulated cell frequencies 0.05, 0.25 or 0.50.

To define the distribution of *N*, we need the column norms of the baseline bulk matrix, which we know in the case of simulated datasets but not real datasets. To infer the baseline bulk matrix, we used an elbow plot to identify the leading principle components of the expression matrix and identified the bulk with the lower principle components. As has been pointed out by others, e.g. [28, 19], this approach gives a biased estimate of the bulk, but an improved estimator lies outside the scope of this work. Figure 3A shows our estimates of the bulk for different cell types and the fit of our RMT approximation. In contrast to the simulated dataset, the RMT approximation fails to capture the top of the bulk’s eigenvalues. Aparicio et al. [2] previously noted this deviation.

**Figure 3.**
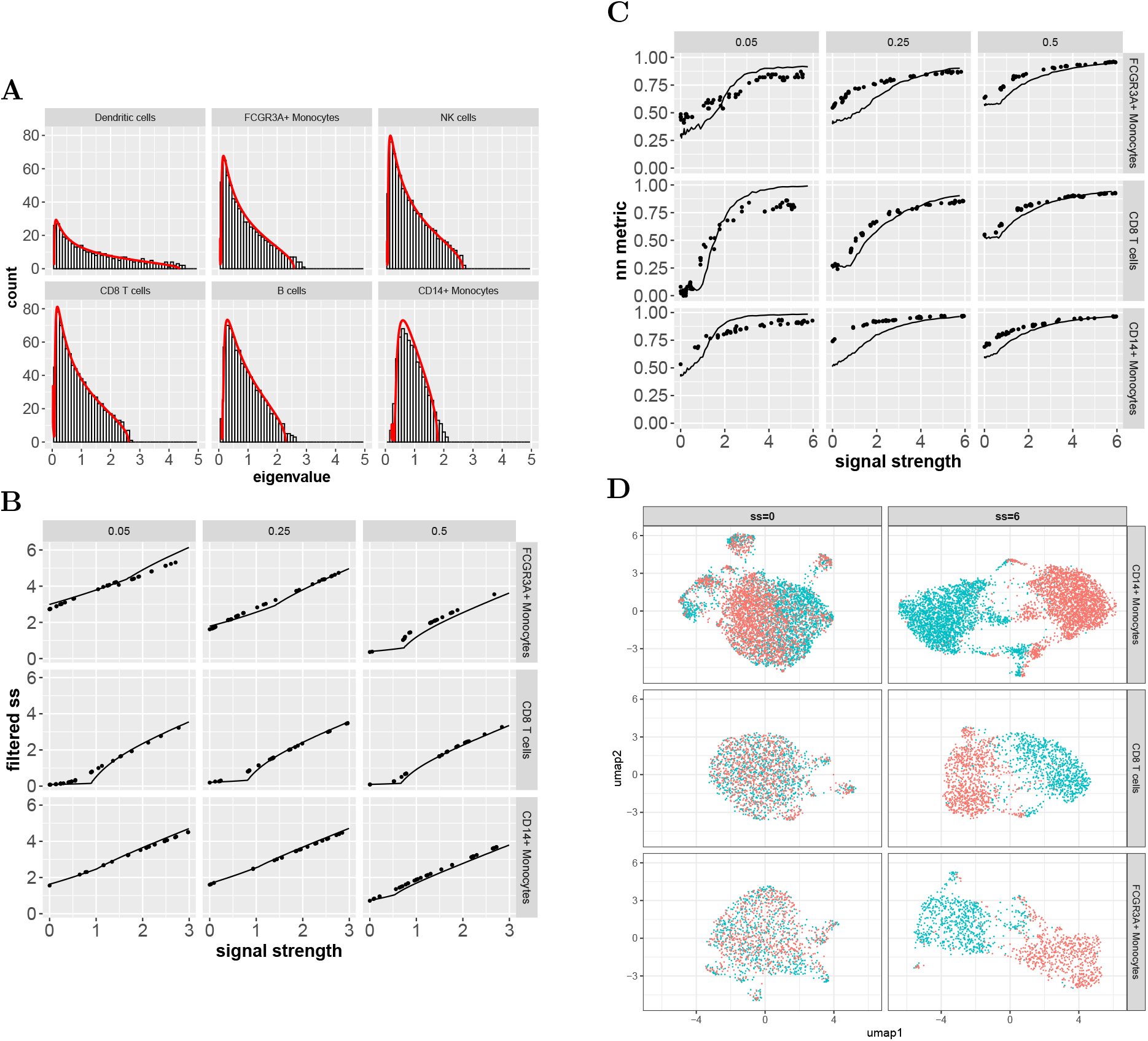
Single Celltype Datasets from Kang et al.. (A) Shown are the estimated bulk (histogram) and the RMT approximation (red line) for the Kang et al dataset restricted to different cell types. The cell types are ordered by sample size, i.e. the number of cells sampled: 428 (DC), 1596 (FCGR3A+ Mono.), 1981 (NK), 2012 (CD8), 2547 (B cells), and 5382 (CD14+ Mono). (B) True (dots) and predicted (lines) values for the filtered signal strength vs signal strength. The columns correspond to different frequencies of IFN stimulated cells. (C) True (dots) and predicted (lines) values for the nearest neighbor metric. (D) UMAPs generated by Seurat for the CD8 T cell and monocyte datasets at signal strengths (ss) of 0 and 6. Colors represent the IFN stimulated (aqua) and control cells (red).

The effect of sample size is not explicit in our formulas, e.g. (6) and (7), but enters through the statistics of *N*. Across the celltypes, as sample size increases, the eigenvalues of the bulk becomes more localized, which affects filtering and clustering. For example, in (7) *b*^2^ represents the maximum eigenvalue of the bulk matrix. As sample size increases across the cell types shown in Figure 3A, *b*^2^ decreases, leading to a smaller noise term under the same signal strength.

Figures 3B,C show the accuracy of our predictions for the filtered signal strength and nearest neighbor metric in the case of FCGR3A+ monocytes, CD8 T cells, and CD14+ monotyes across different frequencies of IFN stimulated cells. The predictions are less accurate than in the simulated dataset due to the errors in approximating the bulk and possibly due to a poorer fit of our statistical model to the perturbation module expression. Nevertheless, the overall trend of both metrics is captured by our predictions.

To see the utility of the signal strength, consider the CD8 T cell and monocyte datasets at a signal strength of 6. Figure 3D shows the UMAPs of these datasets at that signal strength. The UMAPs show similar clustering. Figure 3C shows that all three datasets also have a nearest neighbor metric of roughly 1 at that signal strength. While the signal strengths are the same for the three cell types, the number of ISGs at signal strength 6 is 108, 64 and 49 for the CD8 T cell, FCGR3A+ and CD14 monocyte datasets, respectively. The SNRs of the monocyte ISGs are much higher, leading to a similar signal strength. While our predictions have significant error, the signal strength is providing a measure that captures the overall trend in filtering and clustering levels.

Notably, the filtered signal strength (both true and predicted) in Figure 3B shows a BBP transition for CD8 T cells but not for the monocytes. Indeed, for the CD14+ Monocyte datasets there is a non-zero filtered signal strength when the signal strength is 0 and from that point the filtered signal strength increases roughly linearly. This stark difference reflects the distinction between baseline spikes that are uncorrelated and correlated, respectively, with the perturbation spike.

The baseline spikes in the CD8 T cell dataset are not signficantly correlated with the perturbation module (correlations were less than 0.08), while the baseline spikes of the monocytes are (correlations of greater than 0.30). In the Kang et al datasets, baseline correlation with the perturbation state, defined by IFN stimulation vs control, arises from weak ISGs that we did not designate as perturbation genes. These genes happen preferentially in monocytes. For example, in CD14+ Monocytes, the gene hrasls2 (involved in cell differentiation [39]) is not expressed in control cells but is weakly expressed in 5% of IFN stimulated cells. The resultant correlation of hrasl2 expression with IFN stimulation is 0.023, which is statistically significant but not sufficient to be identified as a ISG under our parameter choices for Seurat’s FindMarker. CD8 T cells have less of these type of genes than monocytes.

The correlation of baseline expression for the monocyte dataset is also reflected in the initial clustering of IFN stimulated cells when the signal strength is 0, as seen through the UMAPs in Figure 3D. At a signal strength of 0, the monocyte datasets (particularly CD14+ monocytes) show clustering of IFN stimulated cells, while the CD8 T cell dataset does not.

### Full Datasets

For the full Kang et al dataset, we consider clustering based on IFN stimulation across multiple cell types. For the other datasets, we consider clustering of a single cell type against a background of multiple cell types. The Mead et al. dataset [27] is sampled from murine, small intestine organoids and includes cells sampled at different time points and conditions. Cell types included 10 different epithelial cells at different stages of differentiation. We set the perturbation genes as genes that were differentially expressed between enteroendocrine cells and all other cell types. In this context, the two perturbation states, *S*_1_ and *S*_2_, were defined by whether a cell was annotated by the authors as enteroendocrine or not. The Ordavas-Montanas dataset [31] is sampled from human nasal biopsies. Using the same approach as for the Mead et al. dataset, we defined perturbation genes to cluster ciliated cell types. The Zheng et al dataset [45] is composed of PBMC and we clustered B cells. See Material and Methods for full details.

Figure 4A shows the fit of *N* to the estimated bulks. The fit has further eroded from the cell type datasets of Kang et al. The fit for the Mead et al. dataset is particularly poor. The poorer fit of the bulks is reflected in less accurate predictions for the filtered signal strength and nearest neighbor metric as shown in Figures 4B,C. However, our predictions still captured the overall trend in the metrics To understand the errors in prediction of the filtered signal, the Ordovas-Montanes et al. dataset is instructive. The failure of *N* to capture the top portion of the bulk leads to an underestimate of the BBP threshold, causing the transition error seen in the Ordovas-Montanes panel of Figure 4B. For the nearest neighbor metric, the Mead et al. dataset is instructive. For small signal strength, our predictions capture the initial increase in the metric, but then overshoot as signal strength increases. Here, the problem is our two state assumption on the perturbation module, not the failure to fit the bulk. Indeed, when we independently permute the entries of each perturbation gene, while not exchanging cells from the two module state *𝒮* 1 and *𝒮* 2, the overshoot disappears as shown in Figure 4C. The UMAPs in Figure 4D correspond to the original and permuted expressions of the Mead dataset at signal strengths of 0 and 5. At signal strength 0, the UMAPs don’t differ, confirming that our permutations don’t change the baseline expression. At signal strength 5, enteroendocrine cells are separated from the other cell types under the permutation but not in the original dataset, corresponding to the higher values in the nearest neighbor metric.

**Figure 4.**
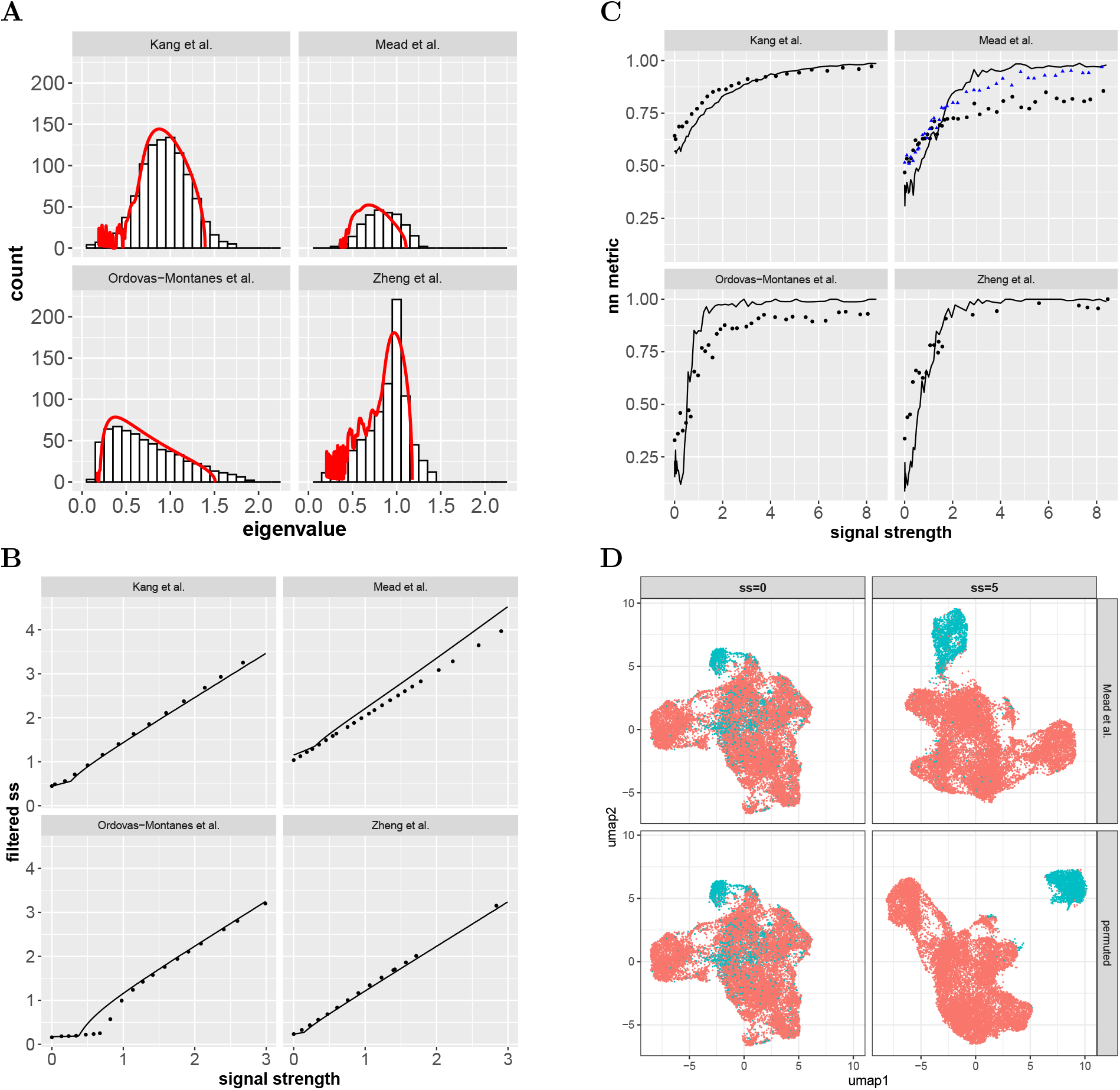
Full Datasets. (A) Shown are the estimated bulk (histogram) and the RMT approximation (red line) for each dataset. (B) True (dots) and predicted (lines) values for the filtered signal strength vs signal strength. (C) True (black dots) and predicted (lines) values for the nearest neighbor metric. The blue dots in the Mead et al panel are generated by permuting the perturbation genes to eliminate all structure not associated with the perturbation states. See text for details. (D) UMAPs generated by Seurat for the unaltered and permuted versions of the Mead et al dataset at signal strength of 0 and 5. Colors correspond to the two perturbation states: enteroendocrine cells (aqua) and other cell types (red).

### Signal Strength of ISG Modules

In application, the nature of the signal strength in different gene modules becomes important. The cell type specificity of the IFN response has been established in a range of studies [41, 38, 23], providing us with a biological context in which to investigate difference in signal strength using the Kang et al dataset. For each cell type in the dataset, we calculated the signal strength of the all the ISGs collectively and the SNR of each ISG. Figure 5 shows the signal strength and SNRs for each cell type. The ISG module of monocytes and dendritic cells has a much greater signal strength than the other cell types, in line with the central role of monocytes and DCs in innate response. Across all cell types, the majority of the signal is carried by roughly 50 ISGs, but no single ISG dominates the signal. The highest SNR, 6.3, occurred in CD14+ Monocytes for the well-known chemokine cxcl10, corresponding to a signal contribution (recall each gene contributes SNR*/*(1 + SNR) to the signal) of 0.86 and composing 2.7% of the signal. Most genes had an SNR of less than 1, reflecting the high level of noise in scRNAseq.

**Figure 5.**
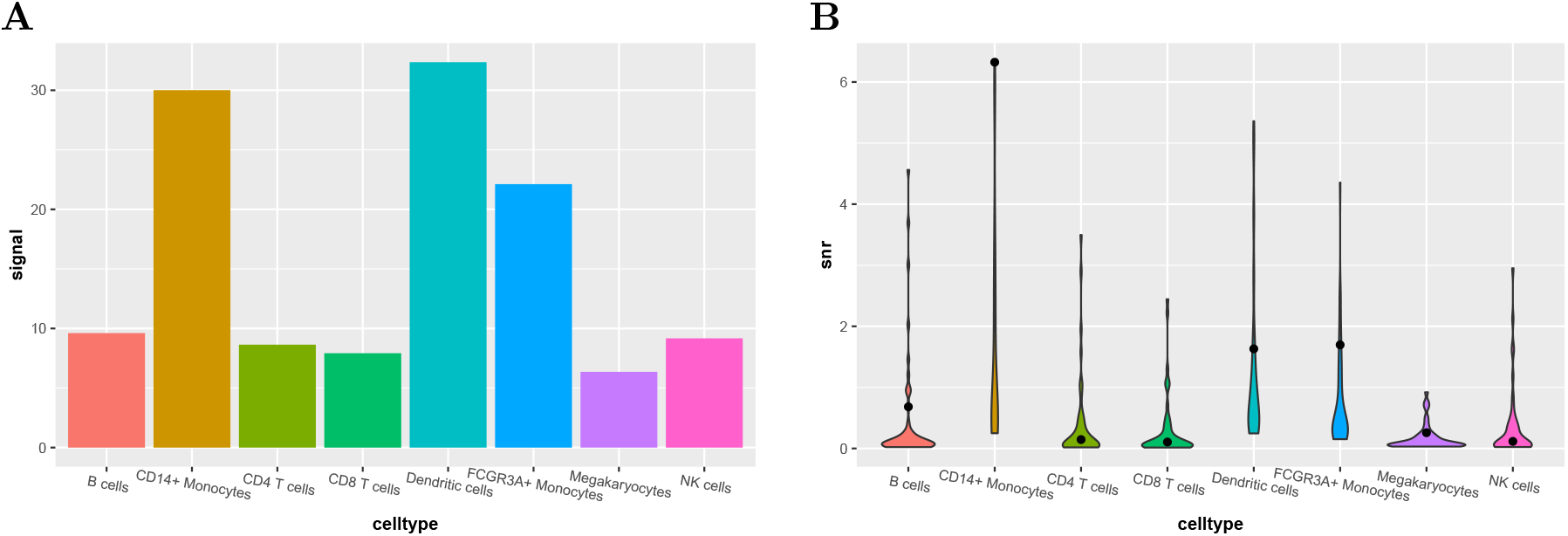
Signal Strength and Signal-to-Noise Ratio of ISG Modules. (A) The signal strength of the ISG module for different cell types in the Kang et al. dataset. (B) The distribution of signal-to-noise (SNR) ratios for each ISG. Black dots show the SNR of cxcl10 for each cell type.

### Effects of Assymetry in the Perturbation State Frequency

Recall that *p*_1_ and *p*_2_ are the fraction of cells in the two perturbation states, *𝒮*_1_ and *𝒮*_2_, respectively, and *p*_1_ + *p*_2_ = 1. These frequencies have a significant impact on the signal strength and SNRs. Notice that the numerator of our formula for SNR, (2), is symmetric in *p*_1_ and *p*_2_ (i.e. exchanging *p*_1_ and *p*_2_ makes no difference), but asymmetric in the denominator when the expression variance in the two states differ. One way this asymmetry may arise is through genes that are not expressed in one cell state, but highly upregulated in the other. A good example is the gene cxcl10. In the CD14+ Monocytes dataset, an average of 64 transcripts are sampled per cell under IFN stimulation, while 0.2 are sampled per cell under the control, making the logFC roughly 6. The variance of cxcl10 expression in the stimulated and control states are 2.1 and 0.8 respectively. If we imagine a population of cells in which 5% of cells are IFN stimulated and 95% control, then cxcl10 will have an SNR of 1.95, while a population with 95% control cells and 5% stimulated will have an SNR of 0.8.

Figure 6A shows the asymmetry in signal strength of the full ISG module when IFN stimulated cells make up 5%, 50% and 95% of the sampled cells. The strongest signal strength occurs at *p*_1_ = *p*_2_ = 0.5 for all cell types, as might be expected. But, perhaps less obvious, the signal strength is stronger when *p*_1_ = 0.05 than *p*_1_ = 0.95 in several of the cell types. Put another way, it may be easier to cluster a small number of IFN stimulated cells on a background of many unstimulated cells than vice versa.

**Figure 6.**
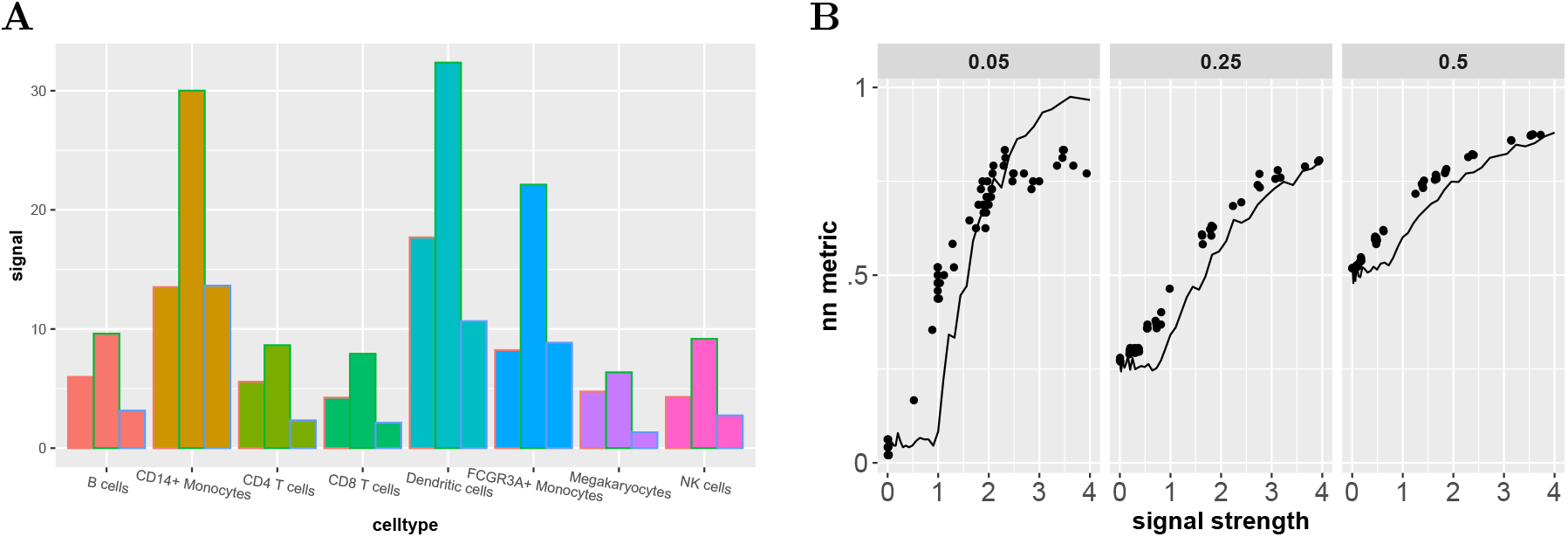
Effects of Assymetry in the Perturbation State Frequencies. (A) For each cell type, the bars from left to right show the module signal strength with *p*_1_ = 0.05, 0.50, 0.95 where *p*_1_ is the fraction of cells that are IFN stimulated. In most cell types, signal strength is higher when *p*_1_ = 0.05 than *p*_1_ = 0.95. (B) True (dots) and predicted (lines) values for the nearest neighbor metric in NK cells in the Kang et al. dataset for the different values of *p*_1_.

Our formulas for the filtered signal strength do not explicitly depend on *p*_1_ and *p*_2_ once the signal strength is fixed. However, for a fixed signal strength, our distance formulas explicitly predict higher nn(*G*) values due to the 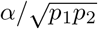 term in (7). As *p*_1_ gets closer to 0, cells of different state will be pushed farther apart in the dimension reduction. Figure 6B shows the nearest neighbor metric for NK cells in the Kang et al dataset over *p*_1_ = 0.05, 0.25, 0.50. The nearest neighbor metric increases much more quickly for *p* = 0.05 than the other frequencies. While the clustering task becomes harder as *p*_1_ becomes smaller, the figure shows roughly equal or greater values for the metric at *p*_1_ = 0.05 versus the other *p*_1_ values. This effect is also seen in the cell types shown in Figure 3C. Importantly as *p*_1_ drops, we expect the signal strength to drop as well, so it is not the case that reducing *p*_1_ leads to better clustering. Rather, over all perturbation modules with equal signal strength, the modules with smaller *p*_1_ will be relatively easier to cluster.

## 3. Materials and Methods

### Datasets

For all real datasets, we started with a raw expression matrix and applied QC to filter the sampled cells. QC was dataset specific and is described below. We then used Seurat’s Find-VariableFeatures function (`selection.method=“vst”,nfeatures=1000`) to select variable genes. For the simulated dataset, QC and gene selection were not relevant. We then used Seurat’s NormalizeData function (`normalization.method=“LogNormalize”`) to generate the normalized matrix. The scaled matrix was created from the normalized matrix by centering the genes and scaling them to length 1, as described in the main text. We used the svdr function in the R irlba package to compute the PCA. We chose 40 pca dimensions except for the Kang et al. and the simulated datasets, for which we chose 20. All UMAP plots were generated using Seurat under the default setting.

For all datasets except Kang et al. and the simulated dataset (see below for details), we selected a particular cell type and used Seurat’s FindMarker function (min.pct=0.05,p_val>0.05) to identify genes differentially expressed in that cell type relative to all other cell types. The perturbation genes were set as the differentially expressed genes and all other genes were base-line genes.

#### Simulated

We constructed a simulated version of the Kang et al. dataset using the splatter R package [43]. We simulated 4 cell types (sample sizes of 500, 1000, 2000 and 5000) and 1060 genes. Each cell type was split equally into IFN stimulated and control labels and the control labels were further split into control-A and control-B types. Genes were split into several categories: 300 genes were not differentially expressed (homogeneous genes), 40 genes per cell type (160 total) were differentially expressed in the particular cell type (cell type genes), 100 genes per cell type (400 total) were differentially expressed in the control-A and control-B samples of the cell type (cell specific ISGs), and 200 genes were differentially expressed in all IFN stimulated cells (joint ISGs). For each gene category, we used the splatter `splatSimulateGroups` function to generate simulated expression. Parameters were set to the default values returned by splatter’s `mockSCE`() function except for the following: `out.prob=0, dropout.type=“none”, lib.loc=10000, lib.scale=0, lib.norm=T, de.facScale=0.15 de.mean=0.1.` The joint ISGs served as perturbation genes while all other genes served as baseline genes.

#### Kang et al

The Kang et al. [20] dataset is composed of 23k PBMC cells in IFN stimulated and control states generated using a 10x Genomics, droplet platform. We downloaded expression files from GEO for accessions GSM2560248 and GSM2560249, as well as the annotation file provided in the GEO entry, which gave cell types, IFN stimulation state, and the Kang et al clusterings. We removed cluster 3, which was noted as a significant outlier that distorts expression processing in [12]. Datasets for individual cell types and cell types with a skewed percentage of IFN stimulated cells were generated by subsampling the full dataset. For each individual cell type, we used Seurat’s FindMarkers function to identify cell type specific ISGs. Perturbation genes for the full dataset were ISGs that were shared across all cell types. Perturbation genes for individual cell types were the cell type specific ISGs.

#### Mead et al

The Mead et al. dataset [27] is composed of 16k cells sampled from murine, intestine organoids and generated using a Seq-Well platform. We downloaded the expression matrix and annotation from the Alexandria project website:

singlecell.broadinstitute.org/single_cell?scpbr=the-alexandria-project We applied QC by restricting to cells with expression in at least 2000 genes, total expression of less than 15000, and less than 20% of expression from mitochondrial genes. Perturbation genes were differentially expressed in the enteroendocrine cell type.

#### Ordavas-Montanas et al

The Ordavas-Montanas dataset [31] is composed of 18*k* cells from human nasal sampling across different levels of inflammation generated using a Seq-Well platform. We downloaded the expression matrix and annotation from the Shalek lab website (`shaleklab.com`). Cell type annotations included 10 cell types including immune and epithelial cells. We applied QC by restricting to cells with expression in at least 2000 genes, total expression of less than 6000, and less than 5% of expression from mitochondrial genes. Perturbation genes were differenetially expressed in ciliated cells.

#### Zheng et al

The Zheng et al dataset [45] is composed of 68*K* PBMC cells generated using a 10x Genomics, droplet based system. We downloaded the expression matrix and annotation, which splits the cells into 11 cell types, from the 10x Genomics github site listed in Zheng et al. We applied QC by restricting to cells with expression in at least 200 genes, total expression of less than 2500, and less than 5% of expression from mitochondrial genes. Peturbation genes were differentially expressed in B cells.

### Details for the RMT Decomposion

#### Notation

Below, we compute expectations and variances, written in the standard way as *E*[] and *V* [] respectively, over the distribution of *X*. For a vector *v*, we write ∥*v*∥ as the Euclidean length of *v*, i.e. the *𝓁*_2_ norm.

The analysis of *Z* is complicated by the library normalization term *L*_*c*_ in (1). In the absence of this term, under our gene module model, each column of *Z* (i.e. each gene) is independent, To derive our theoretical results for *Z*, we replace *L*_*c*_ by *E*[*L*_*c*_] in (1). However, all of our numerical studies are performed on *Z*.

Let *y* be the expression of perturbation gene *g* in the normalized matrix *Y*.

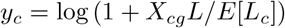

Letting the first *C*_1_ coordinates and the subsequent *C*_2_ coordinates of *y*_*c*_, where *C*_1_ + *C*_2_ = *C*, correspond to cells in state *𝒮*_1_ and *𝒮*_2_, respectively, we can write *v*_*c*_ in block form,

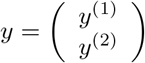

Let 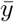 be the mean of the vector *y* and 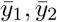 be the means of the two blocks. To scale *y*, and arrive at the corresponding column *z* in *Z*, we center *y* and scale.

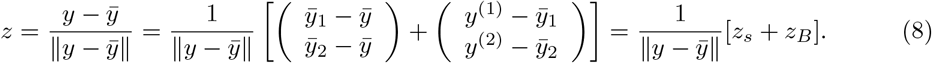

(Above we’ve written 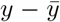 to represent the coordinate wise subtraction of the mean 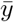 from the vector *y* and similarly for other terms).

The *z*_*B*_ term in (8) will associate with the bulk term since *E*[*z*_*B*_] = 0. To derive the formulas for the spike, we need to relate *z*_*s*_ to *u* and derive a formula for 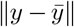. Starting with *u* as 1 for the first *C*_1_ coordinates and 0 for the others, and then centering and rescaling gives,

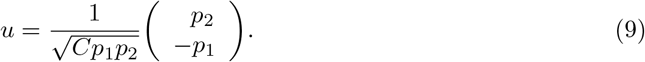

Using 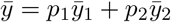

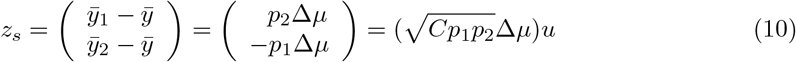

where 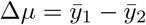. Putting all this into (8),

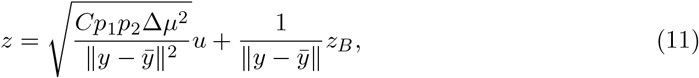

Turning to 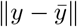, applying a variance decomposition based on the two blocks gives,

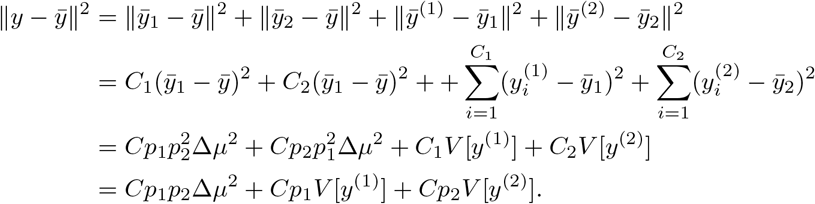

Using the above equation in (11) gives,

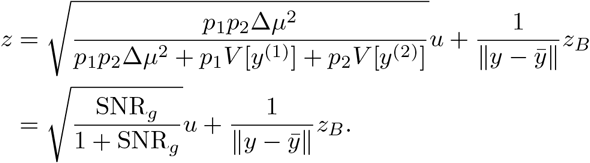

The RMT approximation formulas follow by collecting the columns of *Z*^pert^.

### Analysis of the Signal through the Workflow

Most of the homogenous baseline case is presented in the main text. Here, we just provide more details in deriving the formula for 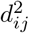. Using (6),

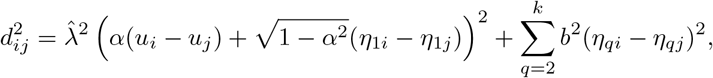

where *u*_*i*_ and *η*_*qi*_ is the *i*th coordinate of *u* and *η*_*q*_, respectively. Since the coordinates of the *η*_*q*_ are normally distributed,

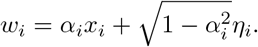

Examining the formula for *u*, (9), shows that *u*_*i*_ − *u*_*j*_ = 0 if cell *i* and *j* are in the same state and 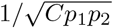 otherwise. Since the factor 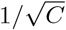 occurs in every term, we drop it without effecting relative distances. We also divide the expression by 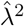. Plugging all this in gives (7)

#### Uncorrelated Baseline

When there is only 1 spike in (4), as for the homogeneous baseline case, RMT results allow us to assume that the non-dominant eigenvectors of 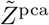, represented by the *η*_*i*_, are normally distributed. However, once baseline spikes are included, the PCA eigenvectors will not typically have normally distributed coordinates. In the uncorrelated baseline case, all spikes are uncorrelated (orthogonal), so we can still apply Theorems 2.9 and 2.10 in [8] to compute 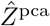. For concreteness, assume that *m < k*. (Recall *m* is the rank of the baseline spike and *k* is the PCA dimension.)

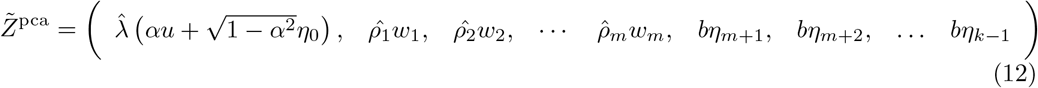

where the *w*_*i*_ for *i* = 1, 2, …, *m* are the PCA-filtered baseline spike signals and have the same form as the filtered perturbation spike signals,

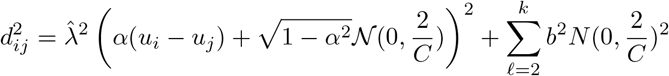

In the case *m > k*, the trailing *bη* terms in (12) are not present. Our estimate for the filtered signal strength, 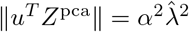 is unchanged from the case of a homogeneous baseline.

The nearest neighbor distance formula (7) must be modified to account for the baseline spikes, but follows from the dimension reduction formula as before.

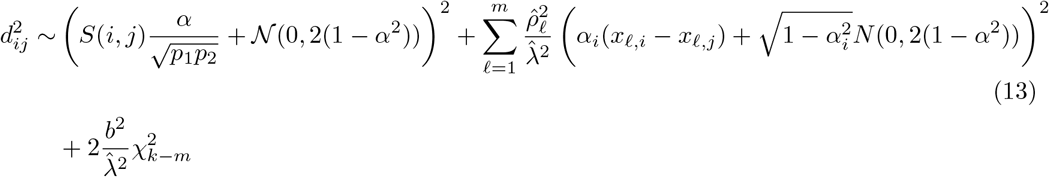

As before, we take 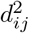 as independent over *j* for fixed *i* and then use a Monte Carlo approach to predict nn(*G*).

#### General Baseline

Finally, we consider a general baseline spike which may be correlated with the perturbation module states. RMT results involving multiple spike terms assume that the spikes are orthogonal (uncorrelated). This is not a restriction since the spike can be orthogonalized by an svd, but in our setting we need to keep track of variation in the *u* direction, so a bit more work is required. To put (5) in orthogonal form, we apply an svd to the spike terms, giving

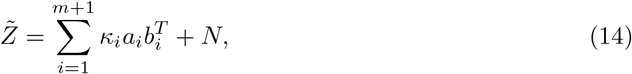

where the *a*_*i*_ and *b*_*i*_ are the singular vectors, which are orthogonal by definition of the svd, and *κ*_*i*_ are the singular values. Importantly, the *a*_*i*_ can be correlated with *u*. To make this correlation explicit, we rewrite *a*_*i*_,

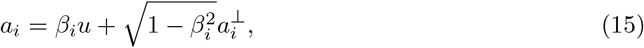

where 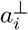*i* is the component of *a*_*i*_ uncorrelated with *u*. From (14), the rest is as before. Each pair *κ*_*i*_, *a*_*i*_ gives a 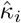 and 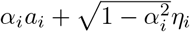. The columns of 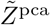 will have the form,

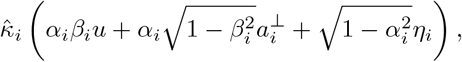

and the filtered signal strength will be given by,

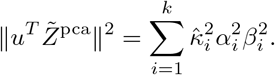

The corresponding distances 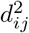 can be computed as before.

### Computation of RMT formulas

The results of Marchenko and Pastur [25] can be applied to *N*, showing that the spectrum of *N*^*T*^ *N* has a well defined limiting density as *C, G* → ∞ with *G/C* fixed. We use the SPEC-TRODE algorithm of Dobriban [13] to compute the limiting density with *G/C* fixed as the ratio of genes to cells in the given scRNAseq dataset.

The matrix *N* fulfills the assumptions of Theorems 2.9 and 2.10 in [8]. Letting *f* (*t*) be the limiting density of *N*^*T*^ *N*, Theorem 2.9 and 2.10 require us to compute integrals involving *f* (*t*). For example, the equation below is equation (4) in [8], which we compute using standard numerical integration.

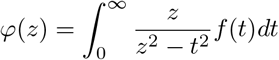

## 4 Discussion

Our results follow a transcriptional signal as it moves through a scRNAseq workflow. We’ve defined a statistical model on the raw expression matrix which assumes that gene expression is characterized by two respective cell states. While other more realistic models are possible, some statistical model of the raw expression matrix is needed to analyze scRNAseq resolution. Otherwise, the relationship between signal and noise cannot be quantified.

Given the raw expression matrix, most workflows normalize the expression in some way. We’ve considered a particular normalization - log normalization - but the particular normalization is not essential to our results. Indeed, our formulas for the gene SNRs, and consequently our formula for signal strength, involve the normalized expression counts, not the raw expression, so the normalized matrix can be seen as the starting point of our analysis. Defining the signal strength from normalized counts reflects the viewpoint that normalization serves to properly define the signal, after known biases have been removed, as has been previously noted [16].

RMT enters into our analysis during the dimension reduction step. Here, our choice of PCA as the dimension reduction method is essential. Non-linear dimension reduction algorithms [24, 40] are commonly used in scRNAseq and it is not clear if RMT can be used in those contexts. Further, our statistical model decomposes the scaled expression matrix into the sum of signal and noise matrices, which allows us to apply particular RMT results. Other statistical models will combine the signal and noise in ways other than a summation, which may or may not be amenable to RMT analysis.

Our results clarify the factors shaping scRNAseq resolution. For a transcriptional signal to lead to clustering, it must make it through the PCA and then sufficiently shape graph construction. Whether a signal makes it through the PCA will depend on the level of noise, which for us is encoded in the bulk matrix. A signal with strength below the bulk’s BBP threshold will be lost, but this will only be true if the signal is not correlated with the other signals in the dataset. Regardless, the signal will pick up noise which may weaken the signal strength. Other factors being equal, more of the signal will pass through the PCA as the bulk becomes more localized. For the datasets we consider, increased sample size associated with a more localized bulk, but our largest dataset was Zheng et al. at 68k cells and whether bigger sample sizes will lead to further localization is unclear.

If a signal passes through the PCA, whether it leads to clustering depends on the other signals in the dataset, which will also be captured in the PCA and will also pick up noise. If the other signals are much stronger, the signal may pass through the PCA but then not lead to clustering. Even in the absence of noise, graph based clustering would still not have infinite resolution because weak signals could be lost against the background of stronger signals. The degree to which this effect shapes current datasets is unclear.

Our analysis has several other implications. Signal strength can be used as a guide for experimental design. For example, consider a particular cell type characterized by differential expression of a particular gene module. The signal strength of the module will depend on the logFC of the genes and the frequency of the cell type within the cell population sampled. If some estimates exist for the logFC, then the frequency needed for clustering of the cell type can be inferred from our formulas for signal strength.

In another direction, we’ve shown that, at least for ISG modules, it can be easier to cluster ISGs on a background of control cells than vice versa. Is this a bias that exists in scRNAseq atlases? Are we missing cell types that turn off genes and preferentially discovering cell types that turn on genes? In the same vein, we’ve shown that the frequency of a cell type matters for clustering. For a fixed signal strength, it is easier to cluster rare cell types than more common cell types. Is this a bias seen in scRNAseq atlases?

Finally, our results provide a framework for comparing differential expression across modules and cell types. The Kang et al dataset is composed of PBMC cell types under IFN stimulation. Signal strength allowed us to compare the ISG modules across cell types with, as expected, monocytes and dendritic cells having the strongest signal. One can imagine application of this approach to non-PBMC. Which cell types can be clustered based on IFN induced, differential expression of their ISGs?

Our results have several limitations. Obviously, we have not been able to consider datasets representing every platform and every cell state. In particular, we have not considered datasets involving cell trajectories. Whether the RMT predictions hold beyond the datasets we consider here will require more work. We have assumed a particularly simple model for a gene module. The accuracy of our predictions may depend on the homogeneity of the module. For example, in the Kang et al datasets, we’ve considered clustering of IFN stimulated cells and in the Zheng et al datasets we’ve considered clustering of B cells. The genes defining these clusters, ISGs for Kang et al and B cell genes for Zheng et al, may form gene modules that conform to our model particularly well. Further work, assuming more complex statistical models for a gene module, is needed. In all our studies, we’ve selected 1000 genes to form the expression matrix. Many researchers use more, e.g 2000 or 3000. As more genes are included, sparsity will increase and our RMT approximations may become less accurate.

For graph construction, we’ve assumed a knn algorithm, while UMAP and shared nearest neighbors are often used. In principle our methods should extend to those algorithms, and we speculate that results will not be very different, but more work is needed. In a similar vein, we have analyzed clustering by considering a metric based on the knn graph. In practice, practitioners often infer discrete clusters by applying the Louvain algorithm [9, 29]. Although it is not clear that metrics based on discrete clustering should serve as a gold-standard for assessing scRNAseq resolution, more work is required to apply our analysis in these contexts.

## Acknowledgements

I thank Brian Rider for helpful discussions regarding RMT.

